# A novel bacterial signal transduction system specifically senses and responds to peptidoglycan damage

**DOI:** 10.1101/2023.05.05.539549

**Authors:** Jianhua Yin, Chaoyi Xu, Xiao Hu, Ting Zhang, Yanqun Liang, Yijuan Sun, Xiangkai Zhen, Yiling Zhu, Yuke Luo, Penshan Shen, Dan Cheng, Yiyang Sun, Jingxiao Cai, Qiu Meng, Tingheng Zhu, Fen Wan, Haichun Gao, Zhiliang Yu

**Affiliations:** College of Biotechnology and Bioengineering, Zhejiang University of Technology, Hangzhou 310014, China; College of Life sciences, Zhejiang University, Hangzhou 310058, China; College of Life sciences, Fujian Normal University, Fuzhou 350117, China; College of Laboratory Medicine, Hangzhou Medical College, Hangzhou 310059, China

**Author notes:** For correspondence (J.Y.); (H.G.); (Z.Y.).

**Keywords:** peptidoglycan, two-component system, antibiotics, signal transduction, response

## Abstract

The peptidoglycan (PG) layer is a mesh-like structure within the cell envelope essential for maintenance of cell shape and resistance to osmotic stress, and therefore is a primary target of many important and widely used antibiotics, such as β-lactams. In Gram-negative bacteria, while signal transduction systems that monitor the state of the inner- and outer-membranes have been extensively studied and well understood, much less is known about how cells sense and respond to PG-targeting stresses. Here we show that a novel bacterial two-component system (PghKR) from *Shewanella oneidensis* is capable of sensing and responding to PG damage. This system is specifically activated in cells exposed to various PG-targeting antibiotics or carrying a defect in PG synthesis, resulting in induced expression of *blaA* and *relV*, which encode a β-lactamase conferring resistance to β-lactams and a small ppGpp synthetase responsible for antibiotic tolerance, respectively. Intriguingly, the PghKR homologs are widespread among several classes of *Proteobacteria* and the periplasmic domain of sensor kinase PghK contains a family 9 carbohydrate-binding module that is required for signal perception, implying that the signals could be the glycan fragments of PG. Overall, our results provide critical insights into the regulation of PG homeostasis in Gram-negative bacteria.

## Introduction

The peptidoglycan (PG) layer, one of the most critical components of the bacterial cell envelope, is necessary for the maintenance of cell morphology and the protection of cells from osmotic rupture (Vollmer et al., 2008; Silhavy et al., 2010). This mesh-like structure is composed of glycan strands built from a repeating disaccharide unit of N-acetylglucosamine-β-1,4-N-acetylmuramic acid (GlcNAc-MurNAc) and short peptides that crosslink glycan strands in proximity. The synthesis of PG begins in the cytoplasm, where the membrane-anchored precursor undecaprenyl-pyrophosphate-linked GlcNAc-MurNAc-pentapeptide (lipid II) is ultimately formed. Then, lipid II is translocated to the outer face of the cytoplasmic membrane by flippase enzymes. In the periplasm, PG glycosyltransferase (PGT) polymerizes the glycan strands, while transpeptidase (TP) forms peptide crosslinks (Typas et al., 2012; Egan et al., 2020). It is well documented that class A penicillin-binding proteins (aPBPs) and SEDS-bPBP protein complexes are responsible for PG synthesis in many rod-shaped bacteria (Meeske et al., 2016; Zhao et al., 2017; Egan et al., 2020; Rohs and Bernhardt, 2021).

There are many antibiotics that target the PG synthetic pathway, and among them, β-lactams represent the oldest and most widely used classes for the treatment of bacterial infections (Lovering et al., 2012; Dörr et al., 2021). This class of antibiotics structurally mimics the D-Ala-D-Ala dipeptide and covalently binds to the active site of TP. As a result, upon exposure, the synthesis of PG is inhibited, leading to PG damage and then cell lysis or death (Kohanski et al., 2010). Other PG-targeting antibiotics, such as vancomycin, moenomycin, and D-cycloserine, cause similar consequences although they interact with different enzymes or function by modifying the PG substrates (Dörr et al., 2021). To survive, bacteria have evolved multiple strategies to be resistant or tolerant to these antibiotics, including the production of enzymes (such as β-lactamase) to hydrolyze or modify antibiotics, activation of stress responses to repair cell damage, and even the stringent response to slow growth as a means to protect cells from lysis (Blairet al., 2015; Dörr et al., 2021).

Bacterial cells employ diverse signal transduction systems to monitor and respond to various environmental stimuli. One of the most common and important systems is the two-component signal transduction systems (TCSs), which are composed of a membrane-associated sensor histidine kinase (HK) and a response regulator (RR) (Laub and Goulian, 2007; Buschiazzo and Trajtenberg, 2019). Upon perception of a specific stimulus, the histidine kinase is autophosphorylated at a conserved histidine and subsequently transfers the phosphoryl group to its cognate response regulator at a conserved aspartate, which binds DNA to modulate gene expression. The TCS involved in Gram-positive bacteria PG metabolism (WalKR) has been extensively studied (Dobihal et al., 2019; Dubrac et al., 2008). However, signal transduction systems that specifically sense and respond to PG damage in Gram-negative bacteria remain poorly understood. To date, the only known example is the VxrAB (also known as WigKR) TCS in *Vibrio cholerae*. VxrAB is activated in response to PG-targeting penicillin and fosfomycin, thereby modulating PG homeostasis and antibiotic tolerance (Dörr et al., 2016; Dörr et al., 2021).

Previous studies have shown that the production of the β-lactamase BlaA in *Shewanella oneidensis*, a representative of ubiquitously distributed γ-proteobacteria, confers natural resistance to β-lactam antibiotics (Fredrickson et al., 2008; Yin et al., 2013; Lemaire et al., 2020). While the expression of *blaA* is conceivably induced by β-lactams (such as ampicillin), it is striking that a similar scenario is found in a strain deficient in the permease AmpG, whose absence results in decreased expression of β-lactamase genes in all other bacteria characterized to date (Yin et al., 2014). More importantly, the expression of *blaA* is constitutively activated in strains lacking the prominent aPBPs (PBP1a, encoded by the *mrcA* gene) and/or its cognate lipoprotein cofactor LpoA, which exhibit severely impaired PG (Yin et al., 2015; 2018; 2020). These observations collectively suggest that expression of *S. oneidensis blaA* is responsive to aberrant PG, and this bacterium could serve as an excellent model to uncover regulatory systems monitoring PG homeostasis (Figure 1A). Moreover, homologs of VxrAB and WalKR have not been found in *S. oneidensis*, although this bacterium possesses 88 TCS proteins (Heidelberg et al., 2002).

**Figure 1.**
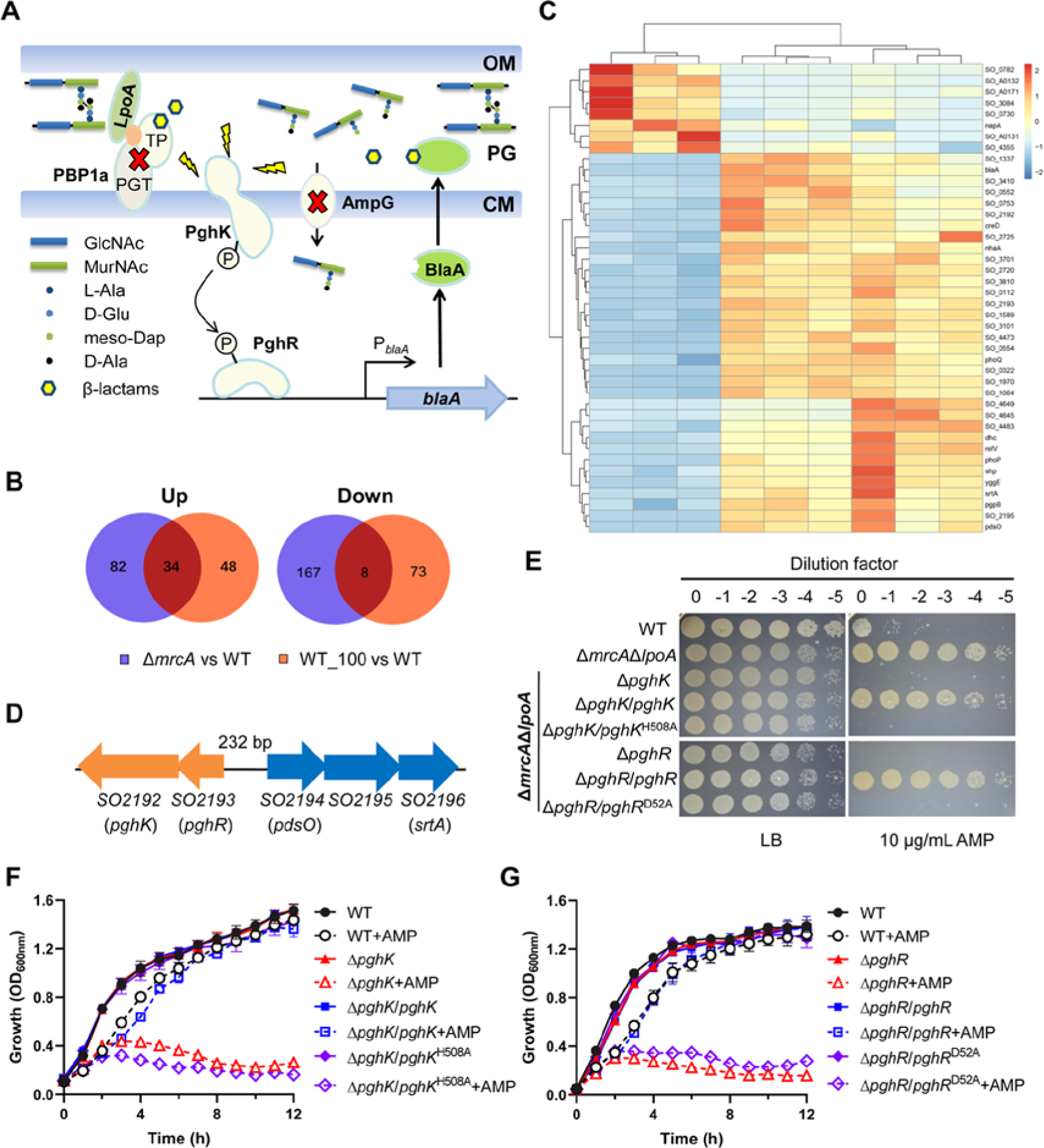
Screening for the regulatory system controlling β-lactamase expression by RNA-seq analysis. **(A)** Schematic of the unknown signal transduction system for the regulation of *blaA* induction in *S. oneidensis*. In contrast to the *ampR*-*ampC* paradigm, deletion of *ampG* resulted in the increased expression of *blaA*, suggesting that *S. oneidensis* possesses a signal transduction pathway for *blaA* induction that is upstream of PG recycling. This pathway involves PBP1a since the inactivation of PBP1a and/or its lipoprotein cofactor LpoA constitutively activated the expression of *blaA*. TP, transpeptidase domain; PGT, glycosyl transferase domain; OM, outer membrane; CM, cytoplasmic membrane. (**B and C)** Venn diagrams (**B**) and Heatmap (**C**) showing the common sets of differentially expressed genes in response to ampicillin treatment (100 μg/mL, AMP_100) and the PBP1a loss (Δ*mrcA*). (**D)** Organization of genes at loci SO2192 to SO2196. The schematic view shows two operons coding for a two-component system (orange) and a marine sortase system (blue). The intergenic region between the two operons is 232 bp in length. The numbers of locus tags and gene names (in parentheses) are given. (**E)** Ampicillin susceptibility assay for *S. oneidensis* strains lacking PBP1a/LpoA complex (Δ*mrcA*Δ*lpoA*) and its derivatives. Cells were grown in LB to an OD_600nm_ ≈ 0.6, 10-fold serial dilutions were prepared, and 3 μL of each dilution was spotted onto LB with or without 10 μg/mL ampicillin (AMP). **(F and G)** Growth of the WT, Δ*pghK* (**F**), Δ*pghR* (**G**) and their derivatives with or without 100 μg/mL ampicillin (AMP). Error bars represent standard deviation from three biological replicates. *pghK*^H508A^ and *pghR*^D52A^ from **(E)-(G)** represent a plasmid expressing PghK variant with the H508A mutation and a plasmid expressing PghR variant with the D52A mutation, respectively.

In this study, by using *blaA* expression as a reporter we identified a previously undescribed TCS, consisting of the sensor kinase PghK and the response regulator PghR, as a central player in sensing and modulating PG homeostasis. PghKR is activated not only in strains defective in PG synthesis but also by exposure to various PG-targeting antibiotics. These findings suggest that the PghKR TCS is capable of monitoring PG damage and triggering a cellular response by subsequently modulating the expression of downstream genes, including those for the stringent response. Intriguingly, the periplasmic domain of PghK contains a carbohydrate-binding module, which is required for signal perception. Overall, our findings reveal a novel signal transduction system responsible for the PG stress response in Gram-negative bacteria.

## Results

### Screening for signal transduction system(s) involved in beta-lactam resistance in *S. oneidensis*

*S. oneidensis* is naturally resistant to β-lactams as a result of the production of the class D β-lactamase BlaA (Yin et al., 2013). Our previous studies demonstrated that the expression of *blaA* is induced by β-lactams (e.g. ampicillin) and constitutively activated in strains lacking PBP1a (Δ*mrcA* and Δ*mrcA*Δ*lpoA*), which are hyperresistant to β-lactams (Yin et al., 2013; Yin et al., 2014; Yin et al., 2015). We hypothesized the presence of the unknown regulator(s) are involved in the regulation of *blaA* expression and that they may also be differentially expressed between cells with *blaA* transcribed at the basal and activated levels. To identify them, RNA-seq experiments were performed to compare the transcriptomes of the wild-type (WT) strain treated with and without 100 μg/mL ampicillin, and the Δ*mrcA* strain. In total, 82 and 81 genes in the WT with ampicillin, and 116 and 175 genes in Δ*mrcA* were significantly up- and downregulated, respectively, when compared with those of the control (untreated WT) (Figure 1B). Interestingly, we found that 34 of the upregulated genes and 8 of the down-regulated genes overlapped in the WT with ampicillin and Δ*mrcA* (Figure 1B and C, Supplementary Table 1). Next, we focused our attention on these overlapping genes.

As we expected, the overlapping upregulated genes included *blaA*, which had ∼14.6- and ∼25.9-fold increases in the WT with ampicillin and Δ*mrcA*, respectively (Figure 1C, Supplementary Table 1). They also included genes encoding proteins implicated in β-lactam resistance or tolerance in other bacteria, such as *creD* and *relV* (Figure 1C, Supplementary Table 1). CreD is a cell membrane integrity protein involved in the CreBC-mediated β-lactam resistance response in *Pseudomonas aeruginosa* (Moya et al., 2009; Zamorano et al., 2014). RelV is a small alarmone synthetase that was found to produce ppGpp in the absence of RelA and SpoT in *Vibrio cholera*, thus contributing to stringent response-mediated antibiotic resistance/tolerance (Das et al., 2009; Kim et al., 2018; Dörr et al., 2021). More importantly, we found that two sets of genes encoding TCSs were significantly upregulated in both the WT with ampicillin and Δ*mrcA* (Figure 1C, Supplementary table 1). One is the *phoPQ* operon, which has been widely reported to play a major role in resistance to antimicrobial cationic peptides and tolerance to β-lactams (Huang et al., 2020; Huang et al., 2021; Murtha et al., 2022). The other consists of *SO2192* and *SO2193*, encoding a histidine kinase and a response regulator, respectively. This TCS is predicted to be involved in controlling a sortase system encoded by an adjacent, divergently transcribed operon, *SO2194*-*SO2196* (Figure 1D). Among them, *SO2194* encodes the sortase system OmpA family protein PdsO, *SO2195* encodes an inter-alpha-trypsin inhibitor family protein, and *SO2196* encodes the marine sortase SrtA. Notably, all three genes were highly upregulated (>17-fold than the WT) in both the WT with ampicillin and Δ*mrcA* (Figure 1C and Supplementary Table 1). This finding coincides with our previous proposal that an unknown TCS may be involved in the regulation of *blaA* expression (Yin et al., 2018). To test this hypothesis, we deleted *phoP*, *phoQ*, *SO2192* and *SO2193* in the background of the Δ*mrcA*Δ*lpoA* strain (Yin et al., 2015). Meanwhile, the *creD* and *relV* genes were also deleted in the Δ*mrcA*Δ*lpoA* strain to test whether CreD and RelV are involved in *blaA* regulation. The spotting assay results showed that two of these mutants, harboring a deletion of *SO2192* or *SO2193*, could not grow on LB agar plates containing ampicillin (Figure 1E and Supplementary Figure 1). Complementation of the two mutants with the WT copy *in trans* restored resistance to ampicillin. Moreover, deletion of *SO2192* and *SO2193* in the WT background also resulted in a significant decrease in ampicillin resistance, which could be fully restored by genetic complementation (Figure 1F and G). However, the ability to grow with ampicillin was not recovered in strains harboring a mutation at the predicted phosphorylation sites (SO2192 with H508A and SO2193 with D52A mutations) (Figure 1E-G). Interestingly, the deletion of *SO2194*, *SO2195* and *SO2196* from either the WT or Δ*mrcA*Δ*lpoA* strain did not affect resistance to ampicillin (Supplementary Figure 1). These results suggest that the SO2192/SO2193 TCS plays an essential role in resistance to β-lactams in *S. oneidensis*, and this role is independent of the sortase system. Based on the studies below, we renamed SO2192 and SO2193 as PghK and PghR for <u>PG h</u>omeostasis sensor <u>k</u>inase and response regulator, respectively.

### PghKR mediates β-lactamase gene expression under conditions of β-lactam exposure and PBP1a loss

To test whether PghK- and PghR-dependent β-lactam resistance is associated with *blaA* expression, the transcription of *blaA* in strains lacking *pghK* or *pghR* in both the samples treated with ampicillin and the untreated control was determined by qRT−PCR (Figure 2A). Upon ampicillin exposure, the transcription of *blaA* in the WT strain was markedly increased, but this response was abolished by the deletion of *pghK* or *pghR*. The complemented strains of Δ*pghK* and Δ*pghR* restored the ability to induce the transcription of *blaA* in the presence of ampicillin. In the background of the Δ*mrcA*Δ*lpoA* strain, *blaA* transcription was highly activated even in the absence of ampicillin (Figure 2B), which is in agreement with our previous results (Yin et al., 2015). Despite this, the removal of *pghK* or *pghR* completely blocked the activation of *blaA* transcription under all tested conditions, and this phenomenon was fully recovered by the expression of the corresponding gene *in trans* (Figure 2B). However, the transcription of *blaA* was no longer induced by ampicillin in mutants harboring PghK^H508A^ or PghR^D52A^.

**Figure 2.**
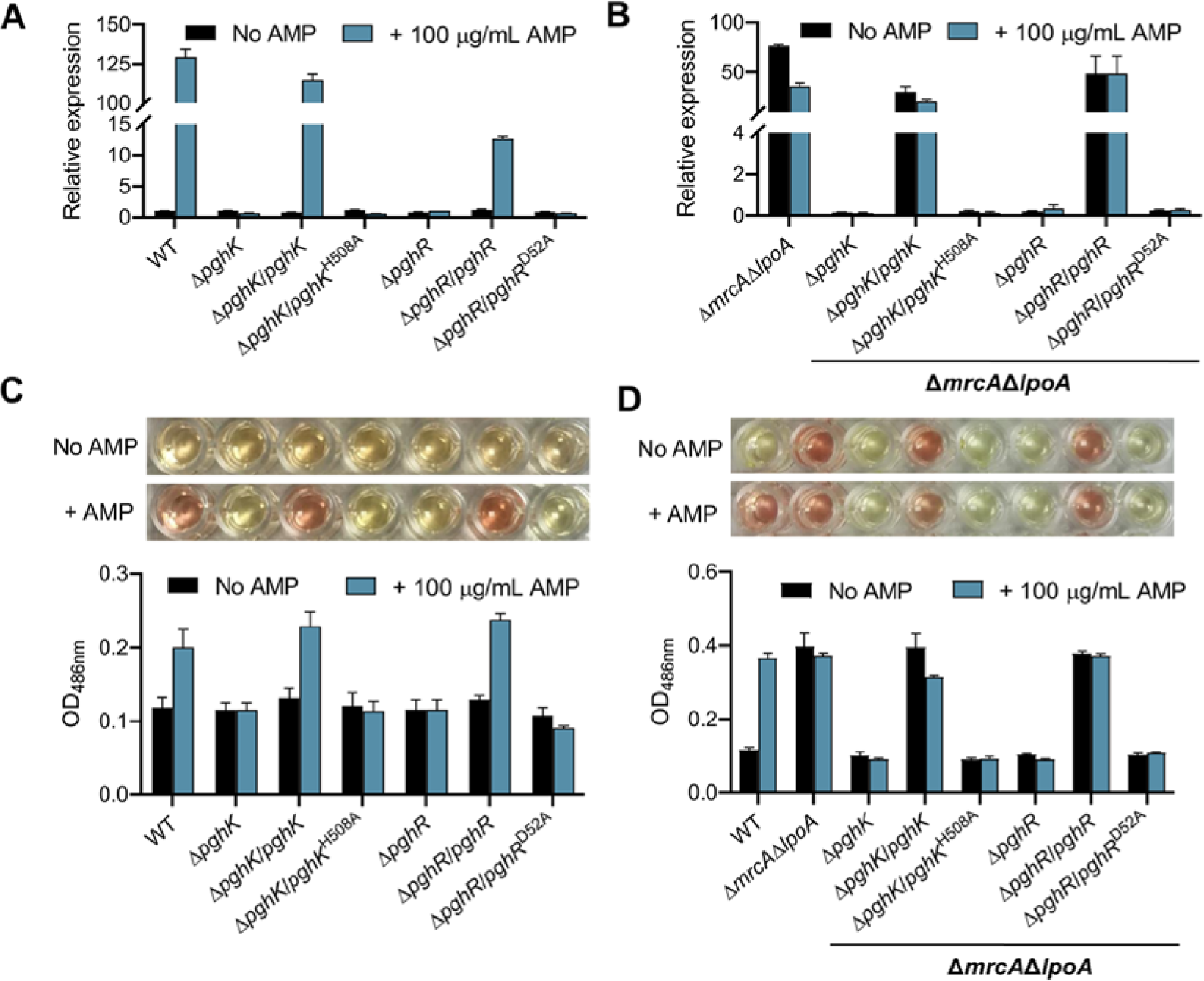
PghKR is required for *blaA* induction in both WT and Δ*mrcA*Δ*lpoA* strains. **(A and B)** Effects of *pghK* and *pghR* deletion on the relative expression of the *blaA* gene in the WT (**A**) and Δ*mrcA*Δ*lpoA* strains (**B**). The *blaA* transcription in the presence or absence of 100 μg/mL ampicillin (AMP) was determined by qRT-PCR analysis. The transcription level of the WT without ampicillin treatment was defined as 1.0. (**C and D**) β-Lactamase activities of the *pghK* and *pghR* mutants in the background of WT (**C**) and Δ*mrcA*Δ*lpoA* strains (**D**). β-Lactamase activity was measured by nitrocefin hydrolysis method. Once hydrolyzed, the yellow substrate nitrocefin (λ max = 390 nm at pH 7.0) is converted to a red product (λ max = 486 nm at pH 7.0) (TOP) and OD_486nm_ was measured (Bottom). *pghK*^H508A^ and *pghR*^D52A^ represent a plasmid expressing PghK variant with the H508A mutation and a plasmid expressing PghR variant with the D52A mutation, respectively. Error bars represent standard deviation from three biological replicates.

We next measured β-lactamase activities directly using the nitrocefin hydrolysis method (Figure 2C and D). Nitrocefin is a chromogenic cephalosporin that can be hydrolyzed by all known β-lactamases, leading to a color change from yellowto red (Li et al., 2016). As we expected, cell lysates from the WT with ampicillin rapidly changed the color of nitrocefin from yellow to red, which exhibited an increase in OD_486nm_ compared with the untreated WT (Figure 2C), supporting that β-lactamase activity is highly elevated by ampicillin. However, strains lacking *pghK* and *pghR* failed to change color upon exposure to ampicillin, but could be restored by expression of wild-type proteins but not phospho-ablative mutations (Figure 2C). Similar results were observed in the background of the Δ*mrcA*Δ*lpoA* strain, which displayed high levels of β-lactamase activity under both basal and induced expression conditions (Figure 2D).

Overall, these results indicate that PghKR is essential for *blaA* expression under conditions of β-lactam induction and PBP1a/LpoA loss.

### PghKR is activated in response to various PG-targeting antibiotics

Given that PghKR is activated not only by β-lactams but also in strains defective in PG synthesis, we hypothesized that PghKR may be activated by other antibiotics targeting PG synthesis. First, we observed the cell morphology in the presence of several antibiotics that target different steps in PG synthesis, including vancomycin (VAN), moenomycin (MmA), and D-cycloserine (DCS) (Figure 3A). The untreated WT cells displayed a typical rod-shaped morphology. Consistent with our previous findings, treatment with ampicillin led to a significant change in cell morphology, which exhibited various aberrant cell shapes including filaments, branches and bulges (Yin et al., 2013). Similarly, aberrant cell shapes were also formed upon exposure to vancomycin, moenomycin and D-cycloserine. Treatment with vancomycin and moenomycin resulted in the formation of spherical cells, while D-cycloserine promoted the formation of filaments and bulges (Figure 3A).

**Figure 3.**
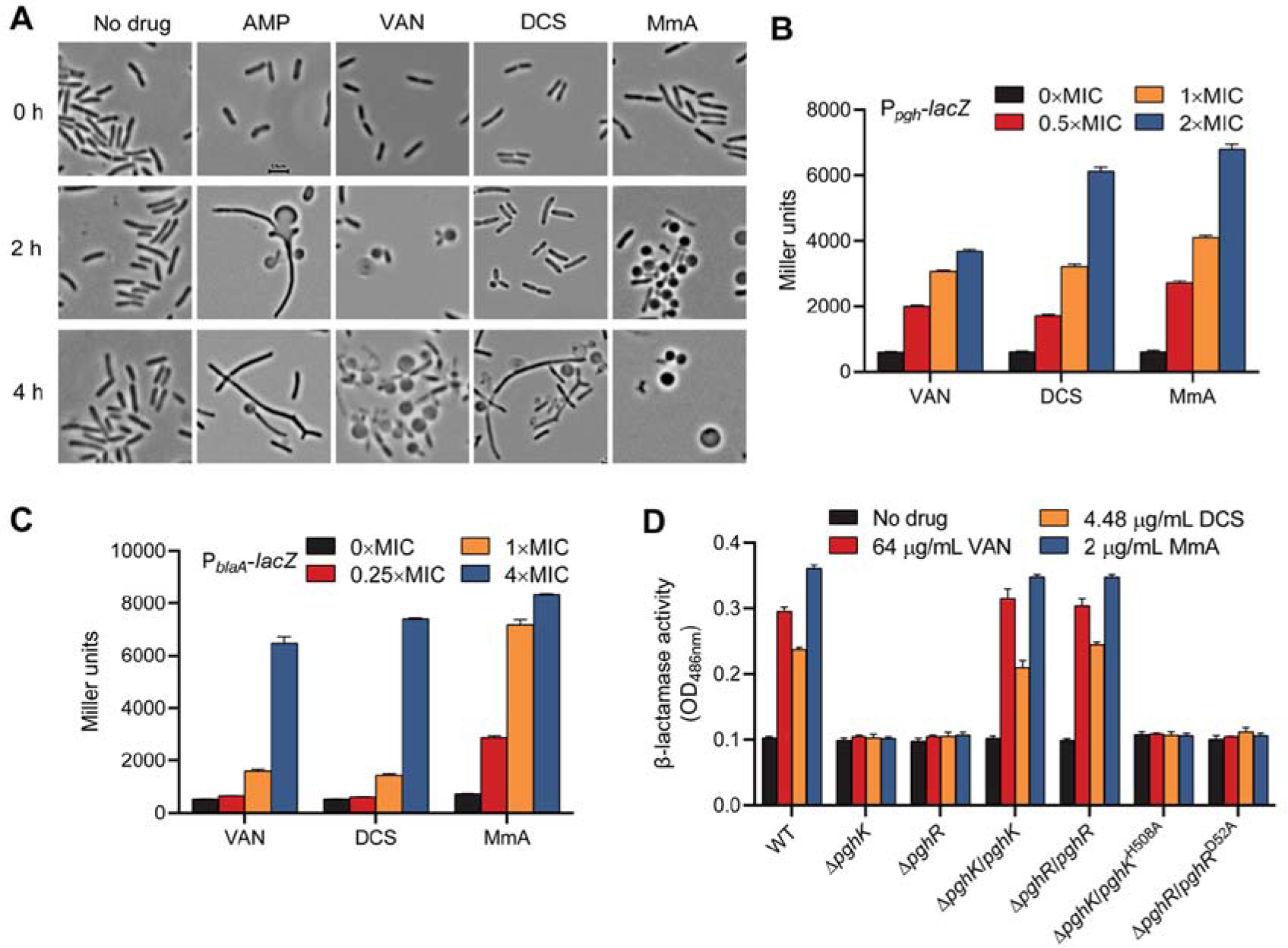
PghKR is responsive to PG-targeting antibiotics. **(A)** Representative Morphology images of the WT following exposure to various PG-targeting antibiotics (100 μg/mL ampicillin [AMP], 64 μg/mL vancomycin [VAN], 9.76 μg/mL D-cycloserine [DCS], and 2 μg/mL moenomycin [MmA]. Cells were grown in LB broth until reaching an OD_600nm_ of ∼0.05, at which point each antibiotic was added. Samples was examined by phase contrast microscopy immediately before and 2 h and 4 h after addition of antibiotics. Scale bar represents 2.5 μm. **(B)** Promoter activity of the *pghRK* operon following exposure to VAN, DCS, and MmA at 0.5 ×, 1 × and 2 × MIC. The MIC values for the WT are listed in Supplementary Table 2. **(C)**, Promoter activity of *blaA* following exposure to VAN, DCS, and MmA at 0.25 ×, 1 × and 4 × MIC. **(D)** Beta-lactamase activity in the WT, *pghK* and *pghR* mutants following exposure to 64 μg/mL VAN, 4.88 μg/mL DCS, and 2 μg/mL MmA, as well as no-antibiotic control. *pghK*^H508A^ and *pghR*^D52A^ represent a plasmid expressing PghK variant with the H508A mutation and a plasmid expressing PghR variant with the D52A mutation, respectively. Error bars represent standard deviation from three biological replicates.

Next, we measured the activities of P*_pgh_* upon exposure to these antibiotics at different concentrations. The results showed that P*_pgh_* was significantly activated upon exposure to vancomycin, moenomycin, and D-cycloserine in a concentration-dependent manner (Figure 3B). Consistently, the activities of P*_blaA_* and β-lactamase also gradually increased in the presence of these antibiotics in a concentration-dependent manner (Figure 3C, Supplementary Figure 2). Moreover, vancomycin and D-cycloserine exposure sufficiently enhanced resistance to ampicillin (Supplementary Figure 2). In the absence of *pghK* and *pghR*, the β-lactamase activity was no longer induced by these antibiotics; however, it could be restored by genetic complementation (Figure 3D). The induced β-lactamase activity was also abolished in strains harboring PghK^H508A^ and PghR^D52A^ (Figure 3D). In contrast to the WT, the addition of vancomycin and D-cycloserine did not enhance the resistance to ampicillin in the *pghK* and *pghR* mutants (Supplementary Figure 2). We also determined the P*_blaA_* activity induced by other classes of antibiotics, including gentamicin, tetracycline, chloramphenicol, erythromycin, polymyxin B and norfloxacin, and the cell membrane-damaging agents SDS and Triton X-100. However, the results showed that none of them were able to enhance P*_blaA_* activity (Supplementary Figure 3).

Collectively, these results strongly suggest that PghKR is specifically activated by PG-targeting antibiotics, which is correlated with PG damage.

### The periplasmic domain of PghK (PghK^SD^) is required for signal transduction

It has been reported that the sensor histidine kinase VbrK in *V. parahaemolyticus* can directly bind β-lactams, leading to the activation of the VbrKR TCS and then the expression of β-lactamase (Li et al., 2016). However, our results clearly showed that PghKR is activated not only by β-lactams but also by structurally diverse PG-targeting antibiotics. Therefore, PghK is unlikely to directly bind β-lactams. Instead, PghK may directly sense PG fragments derived from PG turnover upon exposure to PG-targeting antibiotics. PghK is composed of two transmembrane regions (amino acids 11-28 and 422-443), a periplasmic sensor domain (amino acids 29-421, PghK^SD^) and a cytoplasmic domain (amino acids 444-719) (Figure 4A). Notably, PghK^SD^ is predicted to possess a type 9 carbohydrate-binding module (CBM9, amino acids 45-305) that was initially identified to play an important role in the recognition and hydrolysis of diverse polysaccharides, such as cellulose, chitin and xylan (Guillén et al., 2010). To determine the ligand-binding pocket of PghK, we attempted to predict the structure of PghK^SD^. The best match template returned from Swiss-model analysis is a CBM9 from *Thermotoga maritima* xylanase (PDB access code: 1i8a), but sequence similarity between them is quite low, sharing only 48% coverage (amino acids 50-239) and 19% identity. In contrast, AlphaFold analysis revealed that PghK^SD^ is composed of two subdomains, subdomain I composed of mixed α-helices and β-sheets (34-74, 271-421) and subdomain II composed of multiple β-sheets in parallel (75-270) (Figure 4B). To further explore, we mutated the putative ligand-binding sites predicted by InterProScan search (including H163, S165, I166, R177, Y178, I179, A246, and I289) in the background of the Δ*mrcA*Δ*lpoA* strain. The results showed that single mutations (R177A, Y178A, and I289A) in PghK completely abolished β-lactamase activity regardless of the presence of ampicillin, while other mutations had little effect on β-lactamase activity (Figure 4C, Supplementary Figure 4). Consistently, the Δ*mrcA*Δ*lpoA* strains harboring PghK with single mutations (R177A, Y178A, and I289A) failed to grow on LB plates containing ampicillin (Figure 4D). These results suggest that R177, Y178, and I289 of PghK are essential for the function of PghKR. Interestingly, R177 and Y178 are located in a putative ligand-binding pocket of PghK (Figure 4B), indicating that these residues may be crucial for ligand binding and signal perception.

**Figure 4.**
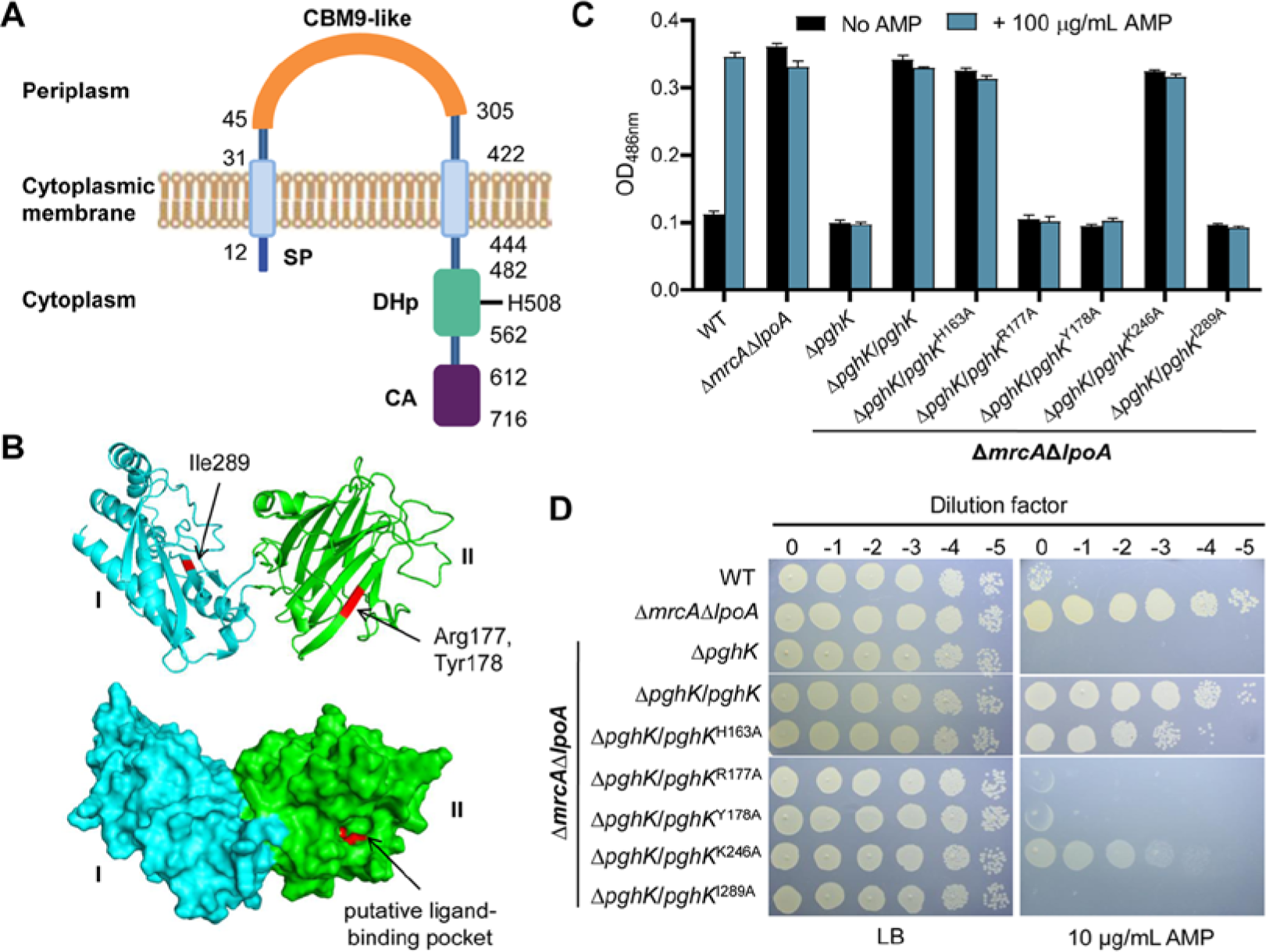
The putative ligand-binding pocket of PghK^SD^ is crucial for signal transduction. **(A)** Domain organization of the histidine kinase PghK. SP, signal peptide; CBM9-like, DOMON-like type 9 carbohydrate binding module; DHp, dimerization and histidine-phosphotransfer domain; CA, catalytic and ATP-binding domain. **(B)** The predicted structure of the periplasmic sensor domain of PghK (PghK^SD^). The structure was obtained by AlphaFold. Upper and lower shows the cartoon and surface image of the structure respectively. The two subdomains of PghK are shown in cyan (I) and green (II), respectively. Three resides (Arg177, Tyr178, and Ile 289) involved in signal recognition are indicated in red. Note that Arg177 and Tyr178 are located in a putative ligand-binding pocket. Error bars represent standard deviation from three biological replicates. **(C)** Beta-lactamase activity in the WT, Δ*mrcA*Δ*lpoA*Δ*pghK* and its derivatives with or without 100 μg/mL AMP. **(D)** Changes in resistance to ampicillin in the WT, Δ*mrcA*Δ*lpoA*Δ*pghK* and its derivatives. Serial dilutions were prepared as in Figure 1 and spotted onto LB with or without 10 μg/mL ampicillin (AMP). *pghK*^H163A^, *pghR*^R177A^, *pghR*^Y178A^, *pghR*^K246A^, and *pghR*^I289A^ from (**C**)-(**D**) represent a plasmid expressing PghK variant with the corresponding mutation. Representative plates from one of three biological replicates are shown.

### PghKR controls ppGpp synthetase responsible for antibiotic tolerance

To explore how PghKR responds to PG damage in addition to β-lactamase induction, the regulon of this TCS was investigated. We focused on the overlapping upregulated genes in both the WT with ampicillin and Δ*mrcA* (Figure 1C, Supplementary Table 1), in which PghKR is activated. The transcription of *relV*, a gene coding for a small ppGpp synthetase that shares 44% identity with *V. cholerae* RelV and contains conserved motifs GYR and EXQX (Supplementary Figure 5), was determined in both Δ*pghK* and Δ*pghR* strains. As expected, the transcription level of *relV* was remarkably increased in the WT upon exposure to D-cycloserine and vancomycin (Figure 5A). However, the induction was nearly abolished in the strain lacking *pghK* or *pghR*, suggesting that the expression of *relV* is largely dependent on the activation of PghKR. Therefore, in addition to the β-lactamase gene *blaA*, PghKR regulon also includes gene coding for ppGpp synthetase RelV.

**Figure 5.**
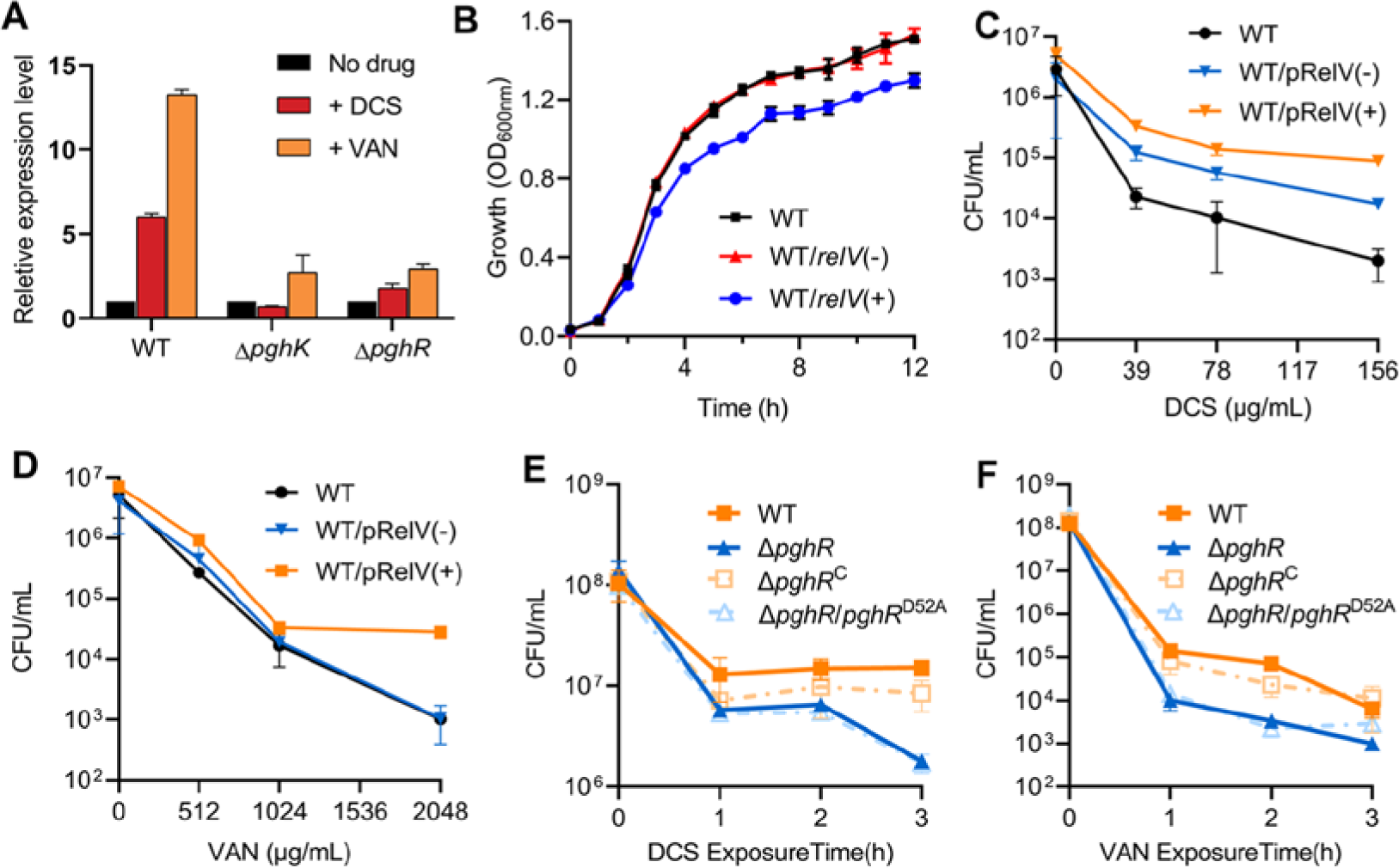
Determination of PghKR regulon. **(A)** The relative expression of the *relV* in the WT, Δ*pghK*, and Δ*pghR* strains upon exposure to 78 μg/mL D-cycloserine [DCS] and 1024 μg/mL vancomycin [VAN]. The transcription level in the WT without antibiotic treatment was defined as 1.0. **(B)** Growth of the *S. oneidensis* WT, WT carrying a plasmid expressing *relV* with [WT/pRelV(+)] or without [WT/pRelV(-)] 0.1% arabinose. **(C and D)** CFU of WT, WT/pRelV(+)] or without [WT/pRelV(-)] 0.1% arabinose after 2 h exposure to DCS (**C**) and VAN (**D**) at different concentrations. **(E and F)** CFU of WT, Δ*pghR*, Δ*pghR* harboring wild-type PghR (Δ*pghR*^C^) and PghR^D52A^ (Δ*pghR*/*pghR*^D52A^) after exposure to 50 μg/mL DCS (**E**) and 1024 μg/mL VAN (**F**) at the indicated time points. Error bars represent standard deviation from three biological replicates.

It has been widely believed that accumulation of ppGpp triggers the stringent response, leading to growth inhibition and then contributing to antibiotic tolerance (Dörr et al., 2021; Irving et al., 2021). To test the effects of *relV* overexpression on cell growth and tolerance to PG-targeting antibiotics, the *relV* gene under the control of the arabinose-inducible promoter P_BAD_ was introduced into the WT strain. As shown in Figure 5B, growth of this strain was evidently inhibited when arabinose was supplemented. More importantly, overexpression of *relV* resulted in more than 10-fold increases in survival after 2 h of exposure to D-cycloserine and vancomycin, respectively, in comparison with the WT strain (Figure 5C and D), suggesting that overexpression of *relV* confers tolerance to PG-targeting stresses. We next determine whether PghKR is involved in antibiotic tolerance. After exposure to D-cycloserine and vancomycin, both Δ*pghK* and Δ*pghR* strains showed ∼10-fold reduction in survival relative to the WT and the respective complementation strains (Figure 5E and F). Similar results were also observed in mutants harboring PghK^H508A^ or PghR^D52A^ (Figure 5E and F).

In conclusion, these results suggest that PghKR controls the overexpression of *relV* in response to PG-targeting antibiotics, which in turn plays an important role in surviving these different antibiotics.

### PghKR homologs are widespread among *Proteobacteria*

Thus far, our data demonstrate that PghKR is a novel TCS that senses and responds to PG-targeted stresses, especially because the sensor PghK contains a CBM9 domain. To ascertain how widespread this signal transduction system might be, the protein sequence of PghK was used as a query to conduct a BLAST search. The results showed that PghK homologs are strongly conserved among several orders of γ-*Proteobacteria* (*Alteromonadales*, *Vibrionales*, *Acidithiobacillales*, and *Beggiatoales* but not in *Enterobacteriales*) (Figure 6A). These proteins are also present in some β-*Proteobacteria* (*Burkholderiales*) and δ-*Proteobacteria* (*Desulfobacterales*) and even in a halophilic archaeon. Intriguingly, Most PghK homologs from β-*Proteobacteria* bacteria lack the CBM9 domain (Figure 6B).

**Figure 6.**
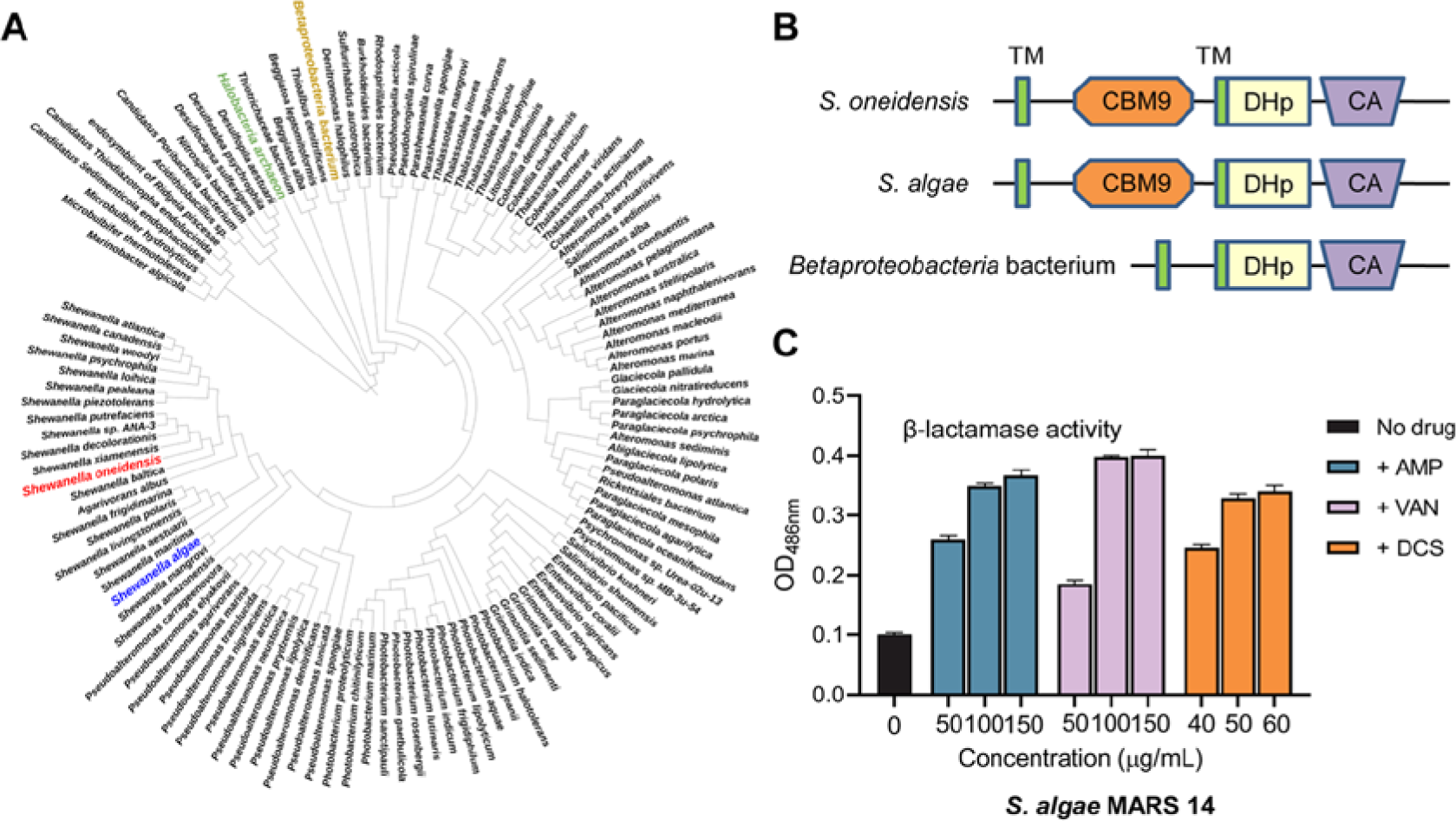
PghKR homologs are conserved among *Proteobacteria*. **(A)** Phylogenetic tree showing distribution of PghK homologs. The amino acid sequence of PghK were used as a query to perform BLAST searches against the UniProt database with an e-value cutoff of 10^−4^ and >32% amino acid sequence identity. The phylogenetic tree was constructed using MEGA and the tree was visualized using iToL (http://itol.embl.de/). PghK homologs in *S. oneidensis* MR-1 (Uniprot ID: Q8EF13), *S. algae* MARS 14 (Uniprot ID: A0A380BZB3), *Betaproteobacteria* bacterium (Uniprot ID: A0A536SEF2) and a Halophilic archaeon (Uniprot ID: A0A7K4E0Y6) were depicted in different colors. **(B)** Domain architectures of PghK homologs mentioned in panel **(A)**. TM, transmembrane domain; CBM9, type 9 carbohydrate binding module; DHp, dimerization and histidine-phosphotransfer domain; CA, catalytic and ATP-binding domain. **(C)** β-lactamase activity in *S. algae* MARS 14 following exposure to AMP (50, 100 and 150 μg/mL), VAN (50, 100 and 150 μg/mL), and DCS (40, 50 and 60 μg/mL), as well as for no-drug control.

To test whether PghKR homologs are also capable of sensing and responding to PG damage, we determined the effects of PG-targeting antibiotics on β-lactamase expression in *S. algae* MARS 14, a multidrug-resistant clinical strain harboring both PghKR and BlaA homologs (Supplementary Table 3). As expected, β-lactamase activities were remarkably induced by ampicillin, vancomycin, and D-cycloserine in a concentration-dependent manner (Figure 6C). Resistance to ampicillin was also improved by vancomycin and D-cycloserine (Supplementary Figure 6). Therefore, these PghKR-harboring bacteria may share a common mechanism for sensing and responding to PG-targeted stresses.

## Discussion

In Gram-negative bacteria, while signal transduction systems that monitor bacterial cell membrane defects have been extensively studied, much less is known about the sensing and response to PG damage (Delhaye et al., 2019). In this study, we show that a previously undescribed TCS in *S. oneidensis*, comprised of histidine kinase PghK and response regulator PghR, is specifically activated in response to various forms of PG damage. Upon exposure to PG-targeting antibiotics and under conditions defective in PG synthesis, the perturbations of PG are sensed by PghK. Similar to other canonical TCSs, PghK transfers the signal to its cognate response regulator PghR, leading to the modulation of gene expression, such as the induction of β-lactamase and ppGpp synthetase to provide either resistance or tolerance to certain PG-targeting antibiotics.

Several TCSs that belong to the cell envelope stress response systems (e.g. CpxAR, BaeSR, RcsCB, and NtrYX) in Gram-negative bacteria have been reported to be associated with PG homeostasis (Delhaye et al., 2016; Delhaye et al., 2019; Mitchell and Silhavy, 2019; Lemmer et al., 2020). However, the genuine signals that these TCSs actually sense are disturbances in the cell membrane rather than PG. TCSs involved in β-lactamase induction, including BlrAB in *Aeromonas* spp. (Alksne and Rasmussen, 1997), CreBC in *P. aeruginosa* (Zamorano et al., 2014), and VbrKR in *Vibrio parahaemolyticus* (Li et al., 2016), are responsive to β-lactams, but the inducible effects of other PG-targeting antibiotics have not yet been described. Our results showed that PghKR in *S. oneidensis* is activated by various PG-targeting antibiotics but not by cell membrane-damaging agents, suggesting that PghKR is a TCS that specifically senses and responds to PG damage. PghKR shares less than 20% sequence identity with all described TCSs, including VxrAB in *V. cholerae*, the only known example for PG damage sensing and response systems in Gram-negative bacteria previously (Dörr et al., 2016).

What signals do PghK directly sense? At present, the signals sensed by PghK and all other TCSs that monitor PG stress (including WalKR in Gram-positive bacteria) remain elusive (Delhaye et al., 2019). It was suggested that the histidine kinase VbrK in *V. parahaemolyticus* can putatively directly bind β-lactams (Li et al., 2016). However, crystal structure analysis showed that the periplasmic sensor domain of VbrK and its homolog VxrA cannot bind to penicillin and related antibiotics directly (Goh et al., 2020; Tan et al., 2021). Recent studies have demonstrated that the cleavage products generated by _D,L_-endopeptidase could serve as the signal for the WalK sensor kinase, although direct evidence is lacking (Dubrac et al., 2008; Dobihal et al., 2019; 2022). In this study, we found that PghKR is responsive to β-lactams, vancomycin, moenomycin, and D-cycloserine. These PG-targeting antibiotics have different molecular structures, excluding the possibility of direct binding of PghK and antibiotics. The signal molecule sensed by PghK may be common PG fragments derived from the action of these PG-targeting antibiotics. PG fragments have been reported to function as signaling molecules to communicate and trigger adaptive responses (Dworkin et al., 2014; Irazoki et al., 2019). Compared with other sensor histidine kinases, PghK is unique since the periplasmic domain of PghK is unusually large (393 amino acids) and contains a family 9 carbohydrate-binding module (CBM9) (Figure 4B). CBM9 was found to be capable of binding insoluble and soluble polysaccharides and a range of soluble oligo-, di- and monosaccharides (Boraston et al., 2001). However, the monosaccharide units of PG, GlcNAc and MurNAc failed to activate PhgKR. We hypothesize that the actual signals sensed by PghK are most likely the glycan fragments of PG. This hypothesis is supported by our previous study that deletion of three lytic transglycosylases (SltY, MltB and MltB2), which cleave the β-(1,4)-glycosidic bond between MurNAc and GlcNAc of the glycan strands, significantly increases β-lactamase expression (Yin et al., 2018).

Both PghKR and VxrAB monitor PG damage, but they respond to PG-targeting antibiotics by distinct mechanisms. PghKR positively regulates the expression of β-lactamase to confer resistance to β-lactams, while VxrAB positively controls genes involved in PG synthesis and negatively controls iron acquisition genes, leading to tolerance to antibiotics (Dörr et al., 2016; Dörr et al., 2021; Shin et al., 2021). In addition to the β-lactamase gene, our results also demonstrate that PghKR modulates the expression of several other genes in response to PG damage, including genes coding for the sortase system and a small ppGpp synthetase RelV. PghKR and its homologs were originally annotated as signal transduction systems controlling the sortase system. Interestingly, sortase enzymes are mainly found in Gram-positive bacteria and function as transpeptidases to anchor surface proteins to the PG, but their roles in Gram-negative bacteria remain unknown (Mazmanian et al., 1999; Spirig et al., 2011). ppGpp is a small nucleotide regulator that induces a stringent response, resulting in the reprogramming of global gene expression and then growth arrest (Irving et al., 2021). Extensive studies have shown that the stringent response is intimately linked to antibiotic resistance and tolerance to various PG-targeting antibiotics due to the regulation of PG synthesis and modulation of ROS production (Kim et al., 2018; Das and Bhadra, 2020; Dörr et al., 2021). Consistently, our results demonstrated that overexpression of *relV* leads to the inhibition of cell growth and tolerance to PG-targeting antibiotics in *S. oneidensis*, finally protecting cells from various PG damage stresses.

To date, PghK is the only known sensor kinase that contains a CBM domain. Bioinformatics analysis revealed that PghK homologs are widespread among several classes of *Proteobacteria*, especially in marine γ-*Proteobacteria* (including the genera *Shewanella*, *Pseudoalteromonas*, *Alteromonas*, *Photobacterium*, *Colwellia*, *Glaciecola* and *Marinobacter*). Remarkably, most PghK homologs from β-*Proteobacteria* lack the CBM9 domain. Because marine environments are rich in polysaccharides (such as chitin, a polymer of GlcNAc), marine bacteria possess a great number of polysaccharide-degrading enzymes and periplasmic solute-binding proteins that contain CBM (Li and Roseman., 2004; Chen et al., 2018). It is likely that the CBM9 domain of PhgK homologs is acquired by horizontal gene transfer from these CBM-containing enzymes. The integration of the CBM9 domain into the sensor kinase in PghK homologs allows bacterial cells to monitor the damage to PG, thus contributing to an adaptive response to a rapidly changing environment.

## Materials and Methods

### Bacterial strains, plasmids and culture conditions

All bacterial strains and plasmids used in this study are listed in Supplementary table 4. All primers used in this study are listed in Supplementary table 5. *Shewanella* and *E. coli* strains were routinely grown in Lysogeny Broth (LB, Difco, Detroit, MI) or Agar (LA) at 37°C and 30°C, respectively. When needed, the medium was supplemented with chemicals at the following concentrations: gentamycin (Gm), 50 μg/mL; kanamycin (Km), 50 μg/mL; ampicillin (AMP), 50 μg/mL; 2,6-diaminopimelic acid (DAP), 0.3 mmol/L.

### In-frame deletion mutagenesis and complementation

In-frame deletion strains for *S. oneidensis* were constructed by the *att*-based fusion PCR method as described previously (Jin et al., 2013). Briefly, two fragments flanking gene of interest were amplified by PCR and then joined by the fusion PCR method. The fusion fragments were introduced into the suicide plasmid pHGM01 by the BP reaction using Gateway BP clonase II enzyme mix (Invitrogen) according to the manufacturer’s instruction. The resulting mutagenesis were maintained in *E. coli* WM3064 (DAP auxotroph) and subsequently transferred into relevant *S. oneidensis* strains via conjugation. Integration of the mutagenesis constructs into the chromosome was selected by resistance to gentamycin and sensitive to sucrose and confirmed by PCR. Verified transconjugants were grown in LB without NaCl and plated on LB supplemented with 10% sucrose. Gentamycin-sensitive and sucrose-resistant colonies were screened and examined by PCR for deletion of the target gene. Finally, the deleted mutants were verified by sequencing.

The promoterless plasmid pHG101 (Wu et al., 2011), the IPTG-inducible expression vector pHGE-P*tac* (Luo et al., 2013) and the arabinose-inducible expression vector pHGC02 (Yin et al., 2015) were used for genetic complementation. For *pghR* and *pghRK* operon, the target gene(s) and its promoter were cloned in pHG101 to generate pHG101-*pghR* and pHG101-*pghRK*. For *pghK*, the target gene was placed downstream of the P*tac* promoter within pHGE-P*tac* to generate pHGE-P*tac*-*pghK*. For *relV*, the target gene was placed downstream of the P_BAD_ promoter within pHGC02 to generate pHGC02-*relV*. All of the resulting vectors were verified by sequencing and then transferred into the corresponding mutant via conjugation.

### Site-directed mutagenesis

pHG101-*pghR* and pHGE-P*tac*-*pghK* were used as the template for site-directed mutagenesis according to the method described before (Sun et al., 2013). Mutated PCR products including *pghR*^D52A^, *pghK*^H163A^, *pghK*^S165A^, *pghK*^I166A^, *pghR*^R177A^, *pghR*^Y178A^, *pghR*^I179A^, *pghR*^K246A^, *pghR*^I289A^ and *pghR*^H508A^ were generated by PCR using primers listed in Supplementary file 5, and then digested by DpnI and transformed into *E. coli* WM3064. After sequencing verification, the resulting vectors were transferred into the corresponding mutants by conjugation.

### RNA extraction, sequencing and analysis

The *S. oneidensis* WT and Δ*mrcA* strains were grown overnight in LB and cultures were diluted 1:100 into 100 mL fresh LB. WT cultures were grown in duplicate. After grown to an OD_600_ of ∼0.05, 100 μg/mL ampicillin was added to one aliquot of WT culture. Cultures were then grown for 2 h and the cells were harvested by centrifugation. Cell pellets were stored at liquid nitrogen for 4 hr. Total RNA was isolated using the RNAiso Plus Kit (TaKaRa, Dalian, China) according to the manufacturer’s instructions. RNA concentration was measured using the NanoDrop ND-1000 Spectrophotometer (NanoDrop Instruments, Wilmington, USA). The removal of rRNA, library construction, RNA sequencing and transcriptome analysis were performed by Zhejiang Tianke High Technology Development Co. Ltd. (Hangzhou, China) as described previously (Hua et al., 2014). RNA sequencing was performed using Illumina HiSeq 2500 platform (Illumina, San Diego, USA). The unique reads were mapped to the genome of *S. oneidensis* (accession numbers NC_004347.2 and NC_004349.1). FPKM (fragments per kilobase per million fragments mapped) of each gene was calculated to compare gene expression levels between the WT and Δ*mrcA*, WT and WT_100 (FDR ≤ 0.05 and absolute value of log_2_ ratio ≥ 1).

### Quantitative real-time PCR (qRT-PCR)

The *S. oneidensis* strains were grown overnight in LB. The cultures were diluted 1:100 into 50 mL LB and were grown to an OD_600_ of ∼0.05. Ampicillin was added to a final concentration of 100 μg/mL. Total RNA was extracted as mentioned above and the cDNA was synthesized using HiScript IIQ Select RT SuperMix (Vazyme, Nanjing, China). qRT-PCR analyses were carried out with the CFX Connect Real-Time PCR Detection System (BioRad, USA) as described previously (Yu et al., 2019). The cycle threshold (*C_T_*) values for each gene were normalized against the *C_T_* values of the 16S rRNA gene.

### Growth of S. oneidensis

Overnight bacterial cultures were diluted 1:100 into 3 mL LB broth and were grown to an OD_600_ of 0.05. Ampicillin or arabinose was added to a final concentration of 100 μg/mL and 0.1%, respectively. Cultures were incubated at 30°C with 200 rpm shaking. The optical density at 600 nm (OD_600_) was recorded every hour after initial inoculation.

### Antibiotic susceptibility assay

Antibiotic susceptibility of *Shewanella* strains was determined both in liquid cultures and on agar plates (Yin et al., 2018). In liquid cultures, the minimal inhibitory concentrations (MICs) for various antibiotics (ampicillin [AMP], vancomycin [VAN], D-cycloserine [DCS], moenomycin [MmA], gentamicin [GEN], norfloxacin [NOR], tetracycline [TET], polymyxin B [PMB], erythromycin [ERY],and chloramphenicol [CHL]) were determined by the CLSI broth microdilution method. On agar plates, the spotting assay was used to assess the susceptibility of *Shewanella* strains to ampicillin. Overnight bacterial cultures were subcultured 1:100 in 3 mL LB and were grown to an OD_600_ of ∼0.6 at 30°C with 200 rpm shaking. Samples were serially diluted and 3 μL of each dilution was spotted onto LB plates, LB plates supplemented with various antibiotics as indicated in the Figure legends. The plates were incubated for 18 h at 30°C and then photographed.

### Promoter activity assay

The promoter activity was determined using a markerless integrative *lacZ* reporter system as described previously (Yin et al., 2014). A fragment containing the target promoter was amplified from genomic DNA and placed in front of the *E. coli lacZ* gene within pHGEI01 (Fu et al., 2014). After sequencing verification, the resultant vector was transferred into relevant *S. oneidensis* strains by conjugation for integration into the degenerated *nrfCD* locus. Overnight bacterial cultures were diluted 1:100 into 3 mL LB broth and were grown to an OD_600_ of 0.05. Ampicillin was added to a final concentration of 100 μg/mL, or other antibiotics were added to a final concentration of 0.25 ×, 1 × and 4 × MIC. After grown for 2 h, cells were harvested by centrifugation. β-Galactosidase activity was determined by measuring color development at 420 nm using a Sunrise Microplate Reader (Tecan), presented as Miller units.

### β-Lactamase activity assay

β-Lactamase activity was determined by nitrocefin hydrolysis method as described previously (Li et al., 2016). Briefly, overnight cultures of the *Shewanella* strains were diluted 1:100 into 3 mL LB broth and were grown to an OD_600_ of ∼0.05. Ampicillin was added to a final concentration of 100 μg/mL, or other antibiotics were added to a final concentration of 0.25 ×, 1 × and 4 × MIC. After grown for 2 h at 30°C with 200 rpm shaking, cells were harvested by centrifugation, washed with PBS, and sonicated. The protein concentration of crude cell extracts was measured using the Bradford method with BSA as a standard (Bio-Rad). The reaction mixture contains 8 μg total proteins and 4 μg nitrocefin (Calbiochem, San Diego, CA). β-Lactamase activity was determined by measuring the absorbance at 486 nm (OD_486nm_).

### Phase-contrast microscopy

Overnight cultures of the *S. oneidensis* WT were diluted 1:100 into 3 mL LB broth and were grown to an OD_600_ of ∼0.05. Various cell wall-targeting antibiotics (100 μg/mL ampicillin, 64 μg/mL vancomycin, 9.76 μg/mL D-cycloserine, and 2 μg/mL moenomycin) were added. After grown for 2 h and 4 h, cells were fixed on a glass slide and visualized with a Motic BA410E phase-contrast microscope. Micrographs were captured with a Moticam Pro 285A digital camera and Motic images advanced 3.2 software (Motic Incorporation, China).

### Antibiotic killing experiments

Overnight cultures of the *S. oneidensis* strains were diluted 1:100 into 3 mL fresh LB broth and were grown to an OD_600_ of ∼0.1. Various concentrations of vancomycin and D-cycloserine were added. After treatment for 1, 2, and 3 h, cells were washed and serially diluted 10-fold in fresh LB broth. 100 μL of each dilution were plated onto LB agar. Viable cell counts (CFU per milliliter) were calculated after 16 of growth.

### Bioinformatics analyses

The domains, predicted phosphorylation sites and putative ligand-binding sites of two-component systems were analyzed using InterPro database (http://www.ebi.ac.uk/interpro/). The structure of the periplasmic sensor domain of SbrK (SbrK^SD^) was predicted using the ColabFold Google Colab notebook “Alphafold _advanced” (Jumper et al., 2021). For construction of phylogenetic tree, the amino acid sequence of PghK were used as a query to perform BLAST searches against the UniProt database with an e-value cutoff of 10^−4^ and >32% amino acid sequence identity. Sequence alignment was performed by ClustalW and the phylogenetic tree was constructed by the Neighbor-joining method in MEGA software. The tree was visualized using iToL (http://itol.embl.de/).

### Statistical analyses

Experimental values are presented as the mean ± standard deviation from three biological replicates. The Student’s *t* test was used to compare statistical differences between the groups of experiment data.

## Acknowledgements

We are grateful to Tobias Dörr for insightful comments on the manuscript. This work was supported by the National Natural Science Foundation of China (Grants 32270044, 31930003 and 31600041), National Key Research and Development Program of China (Grant 2021YFA0909500), Provincial Natural Science Foundation of Zhejiang, China (Grant LY20C010003), and the Fundamental Research Funds for the Provincial Universities of Zhejiang (Grant RF-A2020006).

## Data availability

The RNA sequencing data have been deposited with links to BioProject accession number PRJNA810738 in the NCBI BioProject database (https://www.ncbi.nlm.nih.gov/bioproject/).

## Author Contributions

J.Y., C.X., T.Z., H.G., and Z.Y. designed research; J.Y., C.X., X.H., T.Z., Y.L., Y.-J.S., Y.Z., Y.L., P.S., D.C., Y.-Y.S., and J.C. performed research; X.Z., and Y.Z. contributed new reagents/analytic tools; J.Y., C.X., X.H., T.Z., Y.L., X.Z., Q.M., T.-H.Z., F.W., H.G., and Z.Y. analyzed data; and J.Y., C.X., H.G., and Z.Y. wrote the paper.

The authors declare no conflict of interest.

